# MERIT: A No-Code Platform for Unifying High-Precision Cognitive Assessment Across Laboratory and Real-World Settings

**DOI:** 10.64898/2026.09.13.751236

**Authors:** Stan J. Colcombe, Charles Ay, Jody Brookover, Chris Palmatier, Mike Xiao, Giovanni Salum, Arno Klein, Michael P. Milham

## Abstract

Cognitive tasks are moving from the laboratory onto participants’ own devices. Browser runtimes make that move easy. Their stimulus timing, however, can vary by tens of milliseconds across devices. We asked how much timing error cognitive and neural measures can tolerate. We injected per-trial stimulus-timing jitter into a public EEG dataset (ERP CORE, N = 40) and re-measured three event-related potential components. Under a prespecified criterion of half the between-participant SD, the N170 tolerated jitter up to 15.8 ms, limited by peak-latency bias. The slower MMN and P3 tolerated more than 20 ms. The full jitter distribution predicted the loss. Heavy-tailed jitter cost less than Gaussian jitter of the same RMS. In simulated reaction-time data, the same tail shape determined whether platform error raised or lowered the ex-Gaussian τ, a candidate attentional biomarker. A single RMS timing figure therefore cannot tell whether a platform is adequate for these measures. We built the Mobile Experimental Research Investigative Toolkit (MERIT) to meet these requirements without code. Researchers design tasks in a visual builder that supports reaction-time, continuous-control, drawing, and sequential task families. A Unity engine embedded natively in the Curious iOS and Android apps runs the tasks. Measured with an optical sensor, inter-stimulus-interval SD was 0.71 to 0.96 ms on three current phones and tablets. It was 3.73 ms on a 2019 budget Android phone. Every device measured at least four times below the N170 ceiling. In the laboratory, the same task definition streams event markers to Lab Streaming Layer (LSL) for synchronized EEG and physiological recording. In the field, it runs on ecological momentary assessment schedules. Native mobile players, no-code builders, and EMA platforms each exist as separate systems. MERIT combines them, so the task characterized against brain and body recordings in the laboratory is the same task participants complete at home.

## Introduction

Cognitive psychology and neuroscience have long depended on computerized tasks to probe mental processes, and these tasks demand tight experimental control: stimuli must be presented with precise timing, and responses measured with millisecond precision. Achieving this precision has traditionally required specialized software such as E-Prime or PsychoPy, or custom code written from scratch. At the same time, platforms marketed as accessible to non-programmers, such as the NIH Toolbox or Cambridge Cognition batteries, are typically proprietary and offer little configurability beyond surface parameters. As a result, implementing a new cognitive task can take months of programming or remain infeasible for investigators without coding expertise. This bottleneck slows methodological innovation and constrains who can contribute to it.

Beyond challenges in task development, the field is also grappling with how to extend cognitive assessment into real-world settings. Field and longitudinal studies seek to measure cognition in naturalistic contexts (in homes, schools, clinics, and daily life) to improve ecological validity and capture dynamics that single-session laboratory visits cannot. EMA and burst-sampling paradigms allow repeated cognitive measurement, providing insight into fluctuations associated with context, mood, sleep, and intervention (Trull & Ebner-Priemer, 2013). Deploying cognitive tasks on commodity devices, however, introduces challenges that the field has not fully solved. Web-based experiments delivered through standard browsers can show stimulus presentation deviations of tens of milliseconds across devices, with worst-case deviations reaching approximately 66 ms in some smartphone configurations (Pronk et al., 2020). Variability of this magnitude is acceptable for surveys but undermines reaction-time inference. Bridging the laboratory-to-field gap has therefore generally meant accepting reduced data quality, restricting paradigms to those tolerant of jitter, or commissioning expensive custom application development.

Here, we quantify the practical significance of timing variability on typical outcome measures of brain and behavior. For ERP components measured against task events, jitter injection into a public reference dataset places the tolerable ceiling, under a prespecified materiality criterion of half the between-participant SD, at 15.8 ms for the N170, with slower components tolerating more than 20 ms (see Performance Benchmarking, Figure 2). Browser-class timing variability therefore approaches or exceeds the tolerable limits for fast components rather than sitting safely below them.

A further, less frequently articulated constraint compounds this gap. Laboratory cognitive research increasingly depends on synchronized recordings of brain and body state (EEG, fNIRS, eye-tracking, ECG, electrodermal activity, respiration) acquired alongside task performance. These recordings provide the mechanistic grounding that lets investigators interpret behavioral signatures: what a reaction-time slowing means at the level of cortical excitability, autonomic state, or attentional engagement. When the cognitive instrument used in the laboratory differs from the instrument used in the field, this grounding is broken. Investigators can either characterize the neurophysiological correlates of task performance with high precision in a laboratory setting or sample behavior densely in the field with no direct link back to the underlying physiology, but not both with the same task. A platform that delivers the same instrument in both contexts, with native support for synchronized neurophysiological acquisition in the laboratory and unmodified deployment to participant devices in the field, would help close this gap. Such a platform would allow behavioral signatures observed at home to be anchored to brain and body state patterns established in the laboratory. Closing this loop has been a structural goal of MERIT’s design.

MERIT was designed to meet these three requirements in one system. Investigators author custom tasks without code. A native runtime executes those tasks on participant devices with timing that is measured rather than assumed. The same task definition streams event markers to LSL in the laboratory and runs on scheduled EMA delivery in the field. The platform pairs a visual, no-code task builder with a Unity-based runtime engine embedded in the Curious mobile applications. The remainder of this paper describes the platform’s design rationale, its architecture, the four task families it currently supports, benchmarking results across devices, and representative applications.

### Relation to Existing Platforms

MERIT operates in a crowded space. Code-based desktop systems (E-Prime, PsychoPy, OpenSesame) deliver laboratory timing and hardware synchronization but do not run natively on phones. Browser platforms (Gorilla, jsPsych, PsychoJS, OSWeb) reach every device but inherit the browser timing variability reviewed above. Inquisit is the nearest neighbor. It ships native mobile players with vendor-verified millisecond timing, but tasks are written in its declarative scripting language rather than a visual builder, and physiological synchronization is limited to serial and parallel port signaling on desktop. Presentation is closer to MERIT on the synchronization dimension. Its Presentation Mobile apps run natively on iOS and Android and stream to LSL from the mobile device itself. The vendor has published an end-to-end calibration of that path, measured through LSL against an external event timer (Woods et al., 2017). Tasks, however, must be authored and tested in Presentation for Windows, in its Scenario Description Language and PCL scripting, then packaged and distributed through the vendor’s hosting service. There is no mobile or browser authoring surface, and no scheduling layer for EMA or burst-sampling designs. The mobile timing evidence is a 2017 vendor poster on devices since retired. The vendor describes per-device calibration tooling as still in development. Fixed batteries (NIH Toolbox, CANTAB) offer normed native administration without custom task design.

A different class of system approaches the same problem from the study-management side. MyDataHelps (CareEvolution) runs native iOS and Android apps. Its no-code project builder combines survey delivery, scheduling, wearable ingestion, EHR retrieval, claims data, and remote biosample kitting. It carries large longitudinal cohorts, including the electronic Framingham Heart Study (McManus et al., 2019) and the RECOVER COVID initiative. Its breadth on the health-data side exceeds anything MERIT claims. Cognitive assessment, however, is delivered rather than authored. The available tasks are a fixed set of Apple ResearchKit active tasks, and every cognitive task in that set runs on iOS only. Custom interactive steps are possible, but they require HTML, CSS, and JavaScript inside a webview, which returns the investigator to both coding and browser-runtime timing. No per-device stimulus timing is published, and wearable and clinical data arrive through cloud APIs rather than time-locked streams.

Curious, formerly MindLogger (Klein et al., 2021), provides no-code EMA delivery and is the participant-facing layer MERIT embeds into rather than a competitor. Where LSL appears elsewhere in Table 1 it arrives through desktop plugins, code components, or a scripting-based authoring layer rather than from a no-code task definition. MERIT emits LSL natively from the same no-code task definition, on mobile and desktop alike. A builder-authored task streams into a laboratory’s existing LSL-instrumented hardware, an ecosystem of over 150 device classes (Kothe et al., 2025), without any code. To our knowledge no existing platform combines no-code authoring, a native mobile runtime with measured millisecond precision, native LSL streaming, and EMA or burst-sampling scheduling. The platforms that match MERIT on runtime and synchronization require a scripting language and a Windows desktop to author a task, and provide no scheduling. The platforms that match it on no-code authoring and scheduling run in the browser or deliver a fixed task set. Millisecond precision on a native mobile runtime is shared with two commercial players. MERIT’s contribution is to close both gaps in one system, so that the task characterized against synchronized recordings in the laboratory is the task delivered, unchanged, to participant devices in the field.

**Table 1.**
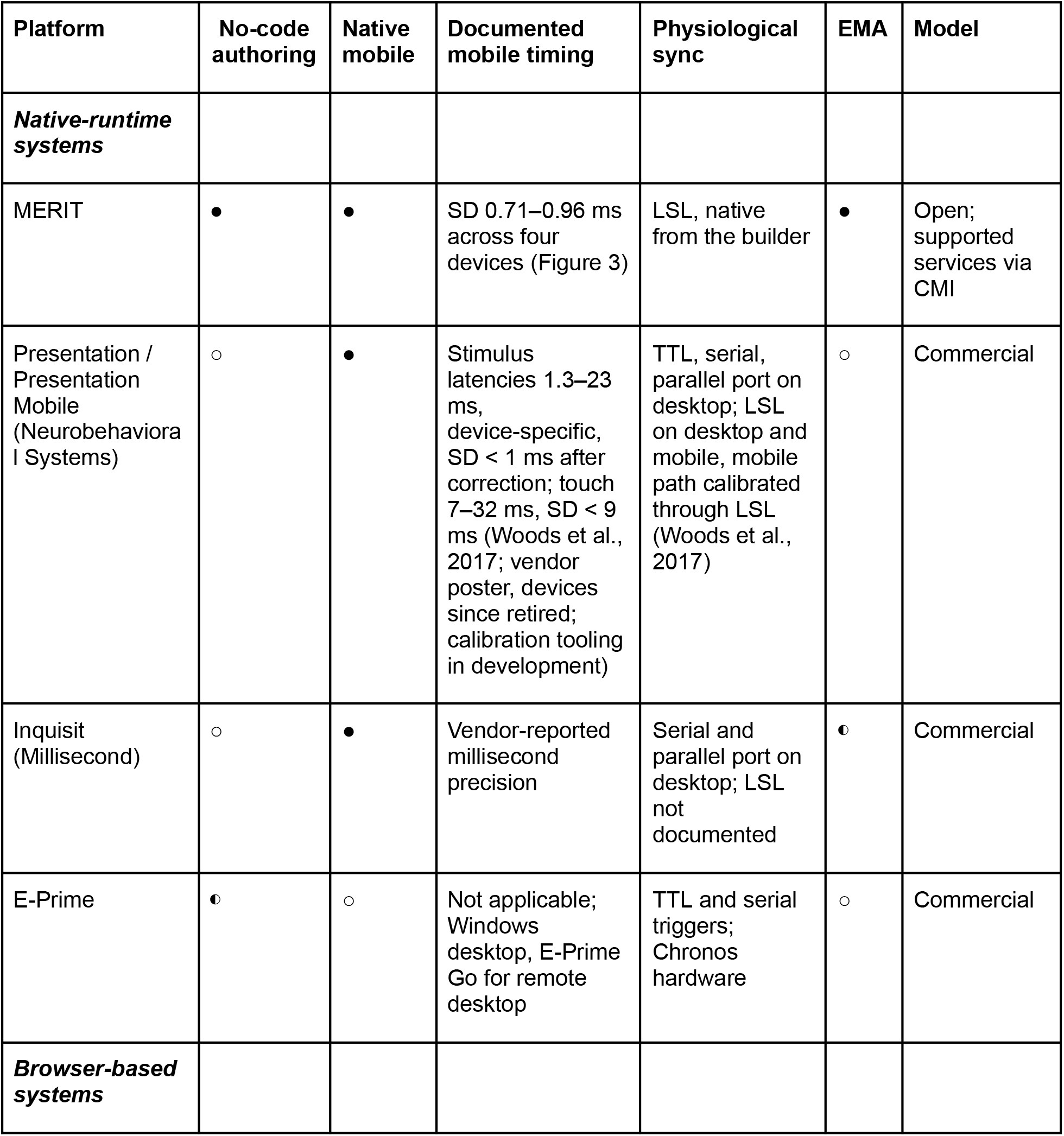
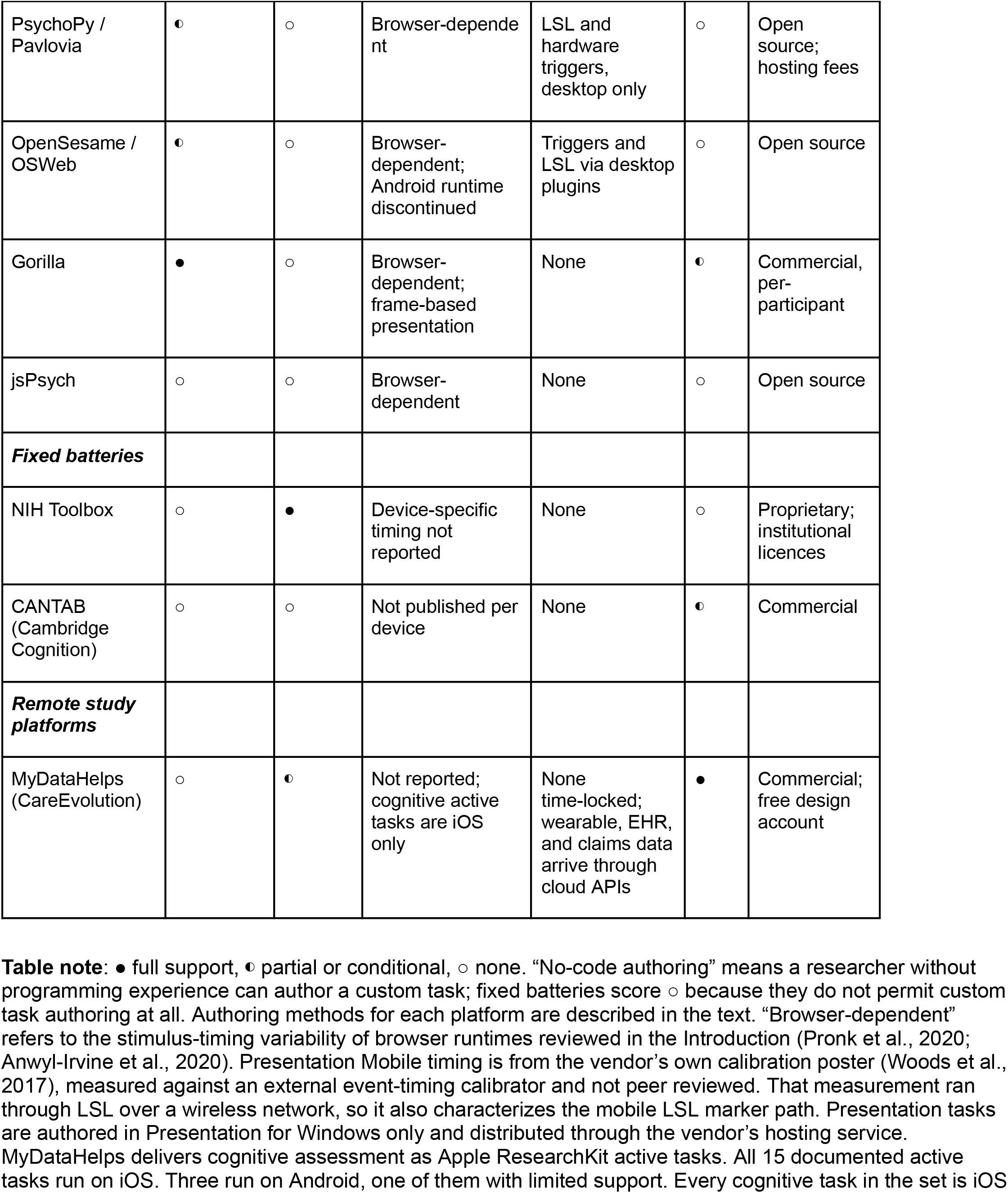
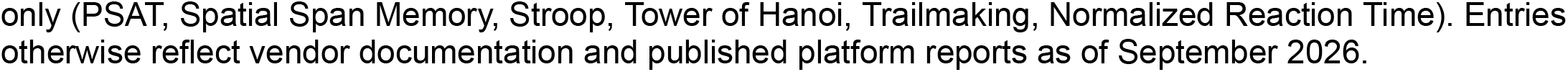
Cognitive task platforms compared on the dimensions MERIT was designed to combine.

#### Platform Overview

MERIT was designed to separate four functions that are often conflated in digital behavioral measurement: task authoring, timing-critical execution, participant-facing study management, and data export. Its architecture consists of a cloud-hosted Builder for creating tasks, a Unity-based Engine for local task execution across platforms, the Curious research management system for consent, scheduling, deployment, and data return (Klein et al., 2021), and a Data Output Manager that supports trial-level exports, real-time LSL streams, raw timing logs, and drawing-task vector traces. This separation allows investigators to create complex cognitive and behavioral paradigms without custom programming, deploy them through longitudinal or EMA-style study schedules, execute tasks locally without network dependence, and synchronize outputs with laboratory or ambulatory data streams. The full system architecture is summarized in Box 1.

#### Box 1. MERIT system architecture

MERIT is organized as a four-part system: a cloud-hosted **Builder** for task authoring, a Unity-based **Engine** for task execution, the **Curious** research management system for participant-facing study delivery, and a **Data Output Manager** for exporting task data into analysis-ready formats.

#### Builder

The Builder is a Unity WebGL application served through a web portal. It allows investigators to design cognitive, behavioral, and sensor-linked tasks without custom programming. The Builder contains five core modules:

- **Resource Manager**. Organizes task assets, including images, audio files, stimulus sets, and drawing templates. Assets are stored in the cloud and pre-loaded before task execution, reducing timing variability caused by on-the-fly loading.
- **Input Manager**. Defines participant responses across deployment contexts, including on-screen touch regions, hardware keys, accelerometer-based motion, and stylus input. Response windows, timeouts, and response rules can be specified at millisecond resolution.
- **Trial Builder**. Provides the main task-authoring surface. Investigators construct trial structures from reusable templates, including simple reaction time, stimulus-mask-response, multi-event sequences, continuous-control paradigms, and drawing-capture tasks. A visual timeline allows researchers to verify the ordering and duration of trial events, including cue onset, stimulus presentation, response windows, and feedback.
- **Block Sequencer**. Organizes trials into blocks and blocks into sessions. Randomization, counterbalancing, conditional branching, adaptive looping, practice blocks, between-subject condition assignment, and multi-day protocol structures can be specified graphically, without custom task logic.
- **Preview mode**. Compiles and runs the current task through the native MERIT Engine rather than through a simulated browser preview. This allows investigators to evaluate rendering behavior and achieved timing during task development, before publishing a task for participant use.

#### Engine

The Engine is the runtime layer. It is written in Unity and embedded as a native module within the Curious iOS and Android applications, which serve as React-based host apps. A thin bridging layer handles platform-specific lifecycle events, input events, and asset delivery. On desktop, the same Unity Engine runs through a corresponding desktop client. This architecture keeps timing-critical task execution within a shared codebase across platforms.

#### Curious integration and participant delivery

Once a task is finalized in the Builder, it is published to the Engine library, versioned, and made available to studies configured in Curious. Participants install the Curious application, consent to enrolled studies, and receive scheduled task assignments. Scheduling supports one-time delivery, repeated sampling at specified intervals, time-of-day windows, and event-contingent triggers suitable for ecological momentary assessment protocols.

#### Local execution and data return

When a task is launched, the Engine downloads the task bundle and executes it locally on the participant device, with no network dependence during task execution. Upon completion, data are returned to the cloud. Data are encrypted at rest by default, both on the participant device before upload and within the cloud storage layer, supporting the security requirements typical of clinical andl research deployments.

#### Data Output Manager

MERIT exports task data in several formats to support different analysis pipelines and integration scenarios:

- **Curious CSV format**. The default export provides one row per trial with stimulus parameters, response timestamps, accuracy, and trial-level metadata. This approach is already well established in the Curious-based questionnaire and EMA deployments, allowing MERIT to integrate seamlessly with existing Curious analysis tooling.
- **Lab Streaming Layer streams**. During task execution, MERIT can emit real-time LSL streams when the participant device is on the same network as an LSL-enabled recording system. Stimulus onsets, response events, and trial markers are timestamped against the LSL clock, enabling synchronization with EEG, eye tracking, physiological monitoring, or other instrumented data streams. LSL is the de facto interchange layer for synchronized neurophysiological recording, with over 150 supported device classes spanning EEG, MEG, fNIRS, eye tracking, motion capture, and peripheral physiology (Kothe et al., 2025). Because support is native, a task built in the Builder streams to a laboratory’s existing LSL-instrumented devices with no additional code. This is the primary integration path for in-laboratory multimodal studies. Inter-stream clock synchronization and offset correction are provided by LSL’s own protocol, validated independently of any client application (Kothe et al., 2025). MERIT’s responsibility is the fidelity of the event markers it hands to that layer. This division of labor, and the end-to-end marker accuracy it implies, is treated under Performance Benchmarking.
- **Raw event logs**. MERIT records frame-level timing information, including per-frame timestamps, requested versus achieved durations, and dropped frame counts. These logs allow investigators to validate timing post hoc and report the achieved precision of a specific deployment.
- **Drawing-task vector traces**. For drawing paradigms, MERIT exports time-stamped coordinate streams suitable for downstream kinematic analysis.

#### Design principle

MERIT separates task authoring, timing-critical execution, participant-facing study management, and data export. This allows investigators to rapidly create and deploy cognitive, behavioral, and sensor-linked paradigms while preserving a common runtime layer for timing-sensitive measurement across mobile, desktop, ecological, and laboratory contexts.

#### Task Families

MERIT currently supports four task families (Figure 1), selected to cover the majority of contemporary cognitive assessment paradigms while exploiting the capabilities of the Unity runtime.

**Figure 1.**
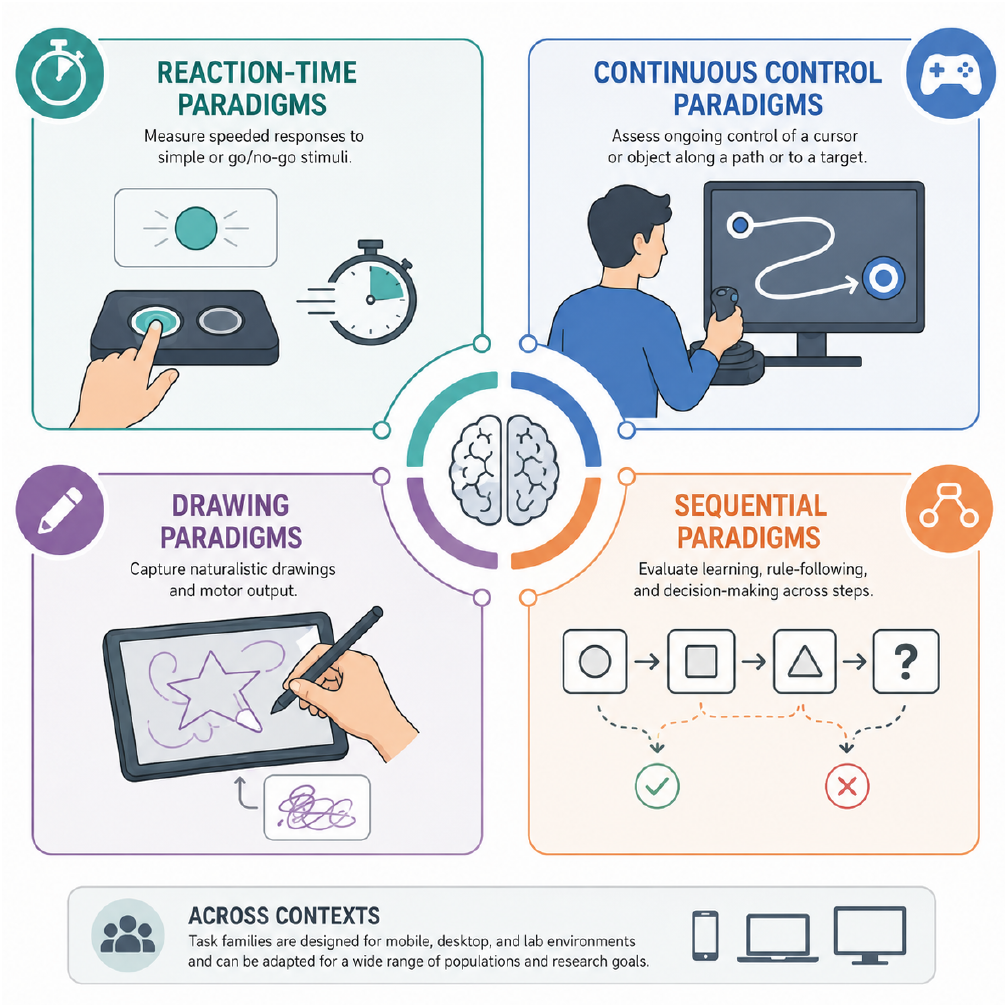
The four MERIT task families. Reaction-time paradigms measure speeded responses to discrete stimuli. Continuous control paradigms measure ongoing control of an unstable element. Drawing paradigms capture stylus or fingertip traces as time-stamped vector streams. Sequential paradigms score ordered actions across a structured stimulus set. All four run on mobile, desktop, and laboratory deployments from the same task definition.

### Reaction-Time Paradigms

The reaction-time family covers classical paradigms in which discrete stimuli prompt discrete responses, with reaction time and accuracy as the primary dependent measures. Tasks like the Flanker, Stroop, n-back, go/no-go, and stop-signal paradigms have all been implemented in MERIT. The psychomotor vigilance task (PVT) is also supported, extending the family to sustained-attention protocols whose outcome measures (lapses and the response-speed distribution) are especially sensitive to platform timing. Every task in the family carries configurable instruction screens and practice blocks with performance gating. Participants can be gated on practice task performance, so that they advance to the main task only after meeting a preset speed and/or accuracy criterion, specified in the Block Sequencer. Item level presentation orders can either be prespecified or pseudorandomized to evenly balance item type presentation orders. These paradigms make the strongest case for MERIT’s timing precision. The inference depends directly on accurate measurement of stimulus onset and response latency, and these are the paradigms for which browser-based deployment has historically been most contentious. Benchmarking results for this family are presented in the next section.

### Continuous Control Paradigms

The continuous control family includes paradigms in which participants exert ongoing motor control to maintain a system within acceptable bounds, with performance characterized by the dynamics of that control rather than discrete trial outcomes. The continuous performance critical stability task (cpCST) is the flagship paradigm in this family and exemplifies the capabilities that the Unity runtime makes possible. In the cpCST, participants control a spatially unstable visual element using the device’s inertial measurement unit (IMU, or ‘accelerometer’). The task algorithmically calibrates each participant’s motor stability threshold (MST) within approximately two minutes and then uses that calibrated threshold to drive a continuous performance phase.

The paradigm requires frame-locked sampling of both the controlled system state and participant input, with no tolerance for timing jitter. cpCST has been validated against established cognitive measures including flanker performance, psychometric IQ estimates, and VO2max (MacKay-Brandt et al., 2025), and has been deployed both in laboratory and remote settings. Continuous control paradigms of this kind are difficult or impossible to implement reliably in browser-based environments and have historically required dedicated laboratory hardware.

### Drawing Paradigms

The drawing family supports digital recapitulation of paper-and-pencil neuropsychological tasks. Investigators upload image underlays and define response capture zones, allowing implementation of tasks such as digit-symbol substitution, Rey-Osterrieth Complex Figure copy and recall, clock drawing, spiral drawing, alphabet writing, and similar paradigms. The same capture engine supports conditional-logic drawing tasks, in which each stroke target depends on the participant’s previous response. Trail Making A and B are the canonical examples, implemented as drawing tasks whose target sequence is governed by the Block Sequencer’s conditional logic. The platform captures stylus or fingertip traces as time-stamped vector streams, preserving kinematic information (velocity, acceleration, pause structure, pen-up and pen-down events) that paper administration discards. Virtually any traditional paper and pencil measure can be digitized via this approach. This enables a class of analyses (e.g., kinematic markers of cognitive slowing or tremor) that are inaccessible to traditional administration, while preserving the underlying paradigm structure that decades of normative data have established.

### Sequential Paradigms

The sequential family covers paradigms in which participants execute an ordered series of actions across a structured stimulus set, with performance characterized by sequence accuracy, completion time, and error patterns. Corsi block-tapping and similar span paradigms are supported. Trail Making, introduced above as a conditional-logic drawing task, is equally a sequential paradigm. Its implementation combines the drawing engine’s trace capture with ordered stimulus presentation, and the engine records per-segment kinematic information by default. Implementing these paradigms benefits from the platform’s combined support for spatial layout, ordered stimulus presentation, and continuous response capture.

These four families have been selected because they cover most contemporary cognitive assessment paradigms while remaining tractable to support at high quality. The platform’s architecture does not preclude additional families (virtual reality navigation, working-memory span tasks with multi-modal stimuli, and others are under active development), but the current release is scoped to paradigms that have been built, tested, and deployed. A planned extension across families is dynamic adaptive testing, in which stimulus difficulty adjusts to the participant’s running performance. Adaptive placement concentrates trials near each participant’s threshold, so a stable estimate arrives in a fraction of the trials a fixed sequence needs. The cpCST’s two-minute calibration of the motor stability threshold is an existing instance of this logic, and the Block Sequencer’s adaptive looping provides the authoring surface for extending it to the other families. Rapid assessment of this kind matters most for EMA and burst-sampling designs, where session length is the binding constraint on sampling density.

#### Performance Benchmarking

Benchmarking requires a target: how much timing error is too much? The answer depends on the temporal structure of the signal being measured, and it can be estimated rather than assumed. We estimated it for the paradigm class most demanding of timing, ERP measurement synchronized to task events through LSL. Per-trial stimulus-timing jitter was injected into a public reference dataset (ERP CORE; Kappenman et al., 2021; N = 40) and each component’s standard estimand was re-measured (Figure 2). We define a materiality threshold as the point at which jitter-induced bias in amplitude or peak latency reaches half the between-participant SD. At that point platform error becomes comparable to the individual differences under study. The resulting ceiling is 15.8 ms of jitter for the N170. The slower MMN and P3 tolerate more than 20 ms. The full bias curves in Figure 2C illustrate the ceiling under alternative criteria. Three properties of this map bear on platform design. First, the loss is predictable in full from the timing-error distribution, because the degraded average equals the clean average convolved with the jitter density (Figure 2A). A platform that measures and reports its own timing distribution therefore lets investigators compute the inferential cost of a deployment rather than guess it. This is why MERIT logs per-frame requested and achieved timing by default.

Second, a single RMS timing figure does not determine the loss. Heavy-tailed jitter, the shape operating-system latency actually takes, destroys less at matched RMS than the Gaussian formula predicts (Figure 2B). The amplitude-only closed form also overstates the N170’s tolerable jitter by 36%, because the binding constraint there is peak-latency bias rather than amplitude. Requirements must therefore be stated against the distribution and the estimand, not a summary statistic. Third, the ceiling sits at the scale of browser-class timing error. Reported browser stimulus deviations of tens of milliseconds (Pronk et al., 2020; Anwyl-Irvine et al., 2020) reach the N170’s 15.8 ms ceiling. Millisecond-level native-engine timing sits an order of magnitude below it. The benchmarks that follow are read against these requirements.

Quantitative validation of timing precision is the empirical foundation of any high-precision task platform, and we conducted benchmarking across the four task families and across representative desktop and mobile devices.

MERIT achieves millisecond inter-stimulus-interval precision on commodity hardware. The feasibility of precision cognitive measurement on consumer devices is a focus of current work, and consequential when it falls short (Pronk et al., 2020). We measured inter-stimulus-interval precision with a Black Box ToolKit optical sensor at 0.25 ms resolution, referenced to a purpose-built USB real-time clock. The SD of timing error was 0.71 ms on an iPhone 16 (2024), 0.84 ms on an iPad (2025), and 0.96 ms on a Galaxy S24 (2024). A Pixel 3a from 2019 showed 3.73 ms. Every interval on every device fell within one 60 Hz frame of its session mean (Figure 3). Read against the requirements in Figure 2, the three current devices sit more than an order of magnitude below the N170’s 15.8 ms ceiling. The six-year-old Pixel 3a sits a factor of four below it. At 1 ms of jitter the N170 retains more than 99.8% of its amplitude, and even at the Pixel’s 3.73 ms it retains 98%. On the axes of Figure 4A, 3.73 ms inflates a 50 ms within-person SD by 0.28%. These measurements locate the open problem elsewhere. Single-stream timing is no longer the central limiting factor for on-device cognitive measurement. What remains is cross-stream alignment: locking a task event on one device to a physiological or neural event on another, independently clocked device (Plant & Turner, 2009).

**Figure 2.**
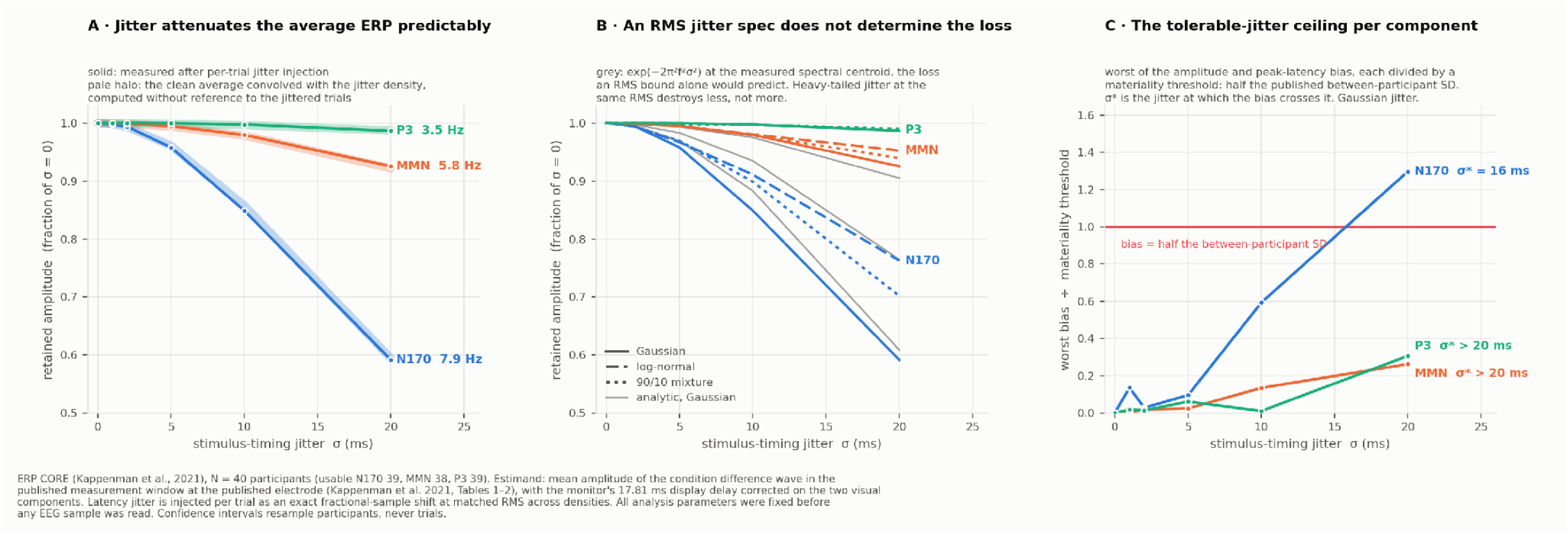
Timing precision required for ERP-synchronized cognitive assessment. We injected per-trial stimulus-timing jitter into a public reference dataset and re-measured each component’s standard estimand (ERP CORE; Kappenman et al., 2021; N = 40; usable N170 39, MMN 38, P3 39). The estimand is the mean amplitude of the condition difference wave in the published measurement window at the published electrode. The recording monitor’s 17.81 ms display delay was corrected on the two visual components. (A) Retained amplitude under Gaussian jitter of SD σ, one line per component, ordered by each component’s measured spectral centroid. The pale halo shows the clean average convolved with the jitter density. That prediction uses no information from the jittered trials, so the degraded average is predictable in full from the timing-error distribution. (B) The same sweep under two heavy-tailed densities at matched RMS, a log-normal and a 90/10 Gaussian-exponential mixture. Grey curves show the analytic Gaussian attenuation exp(−2π^2^f^2^σ^2^) at the measured spectral centroid. Densities with identical RMS produce different losses, and heavy-tailed jitter destroys less than the Gaussian form predicts. An RMS timing specification alone therefore does not determine the inferential cost. (C) The worse of the amplitude and peak-latency bias, each divided by a materiality threshold of half the published between-participant SD. σ* is the jitter at which the bias crosses that threshold. The ceiling is 15.8 ms for the N170, bound by peak latency rather than amplitude. Jitter was injected as exact fractional-sample shifts. All analysis parameters were fixed before any EEG data were read. Confidence intervals resample participants, never trials.

**Figure 3.**
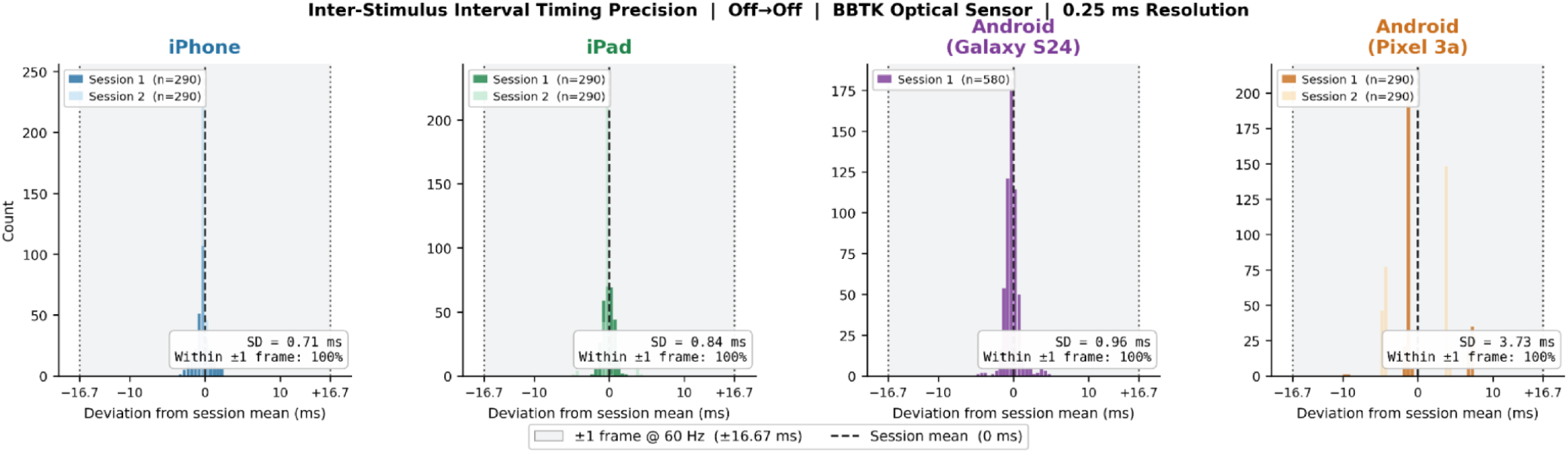
Measured timing precision of MERIT on iOS and Android devices. Inter-stimulus-interval precision, measured off-to-off with a Black Box ToolKit optical sensor at 0.25 ms resolution. Histograms show each interval’s deviation from its session mean. The iPhone 16, iPad, and Pixel 3a were each measured over two sessions of n = 290 intervals. The Galaxy S24 was measured over one session of n = 580. The shaded band marks one 60 Hz frame (±16.67 ms) either side of the session mean. The SD of timing error is 0.71 ms on the iPhone 16 (2024), 0.84 ms on the iPad (2025), 0.96 ms on the Galaxy S24 (2024), and 3.73 ms on the Pixel 3a (2019). Every interval on every device falls within one frame of the session mean.

**Figure 4.**
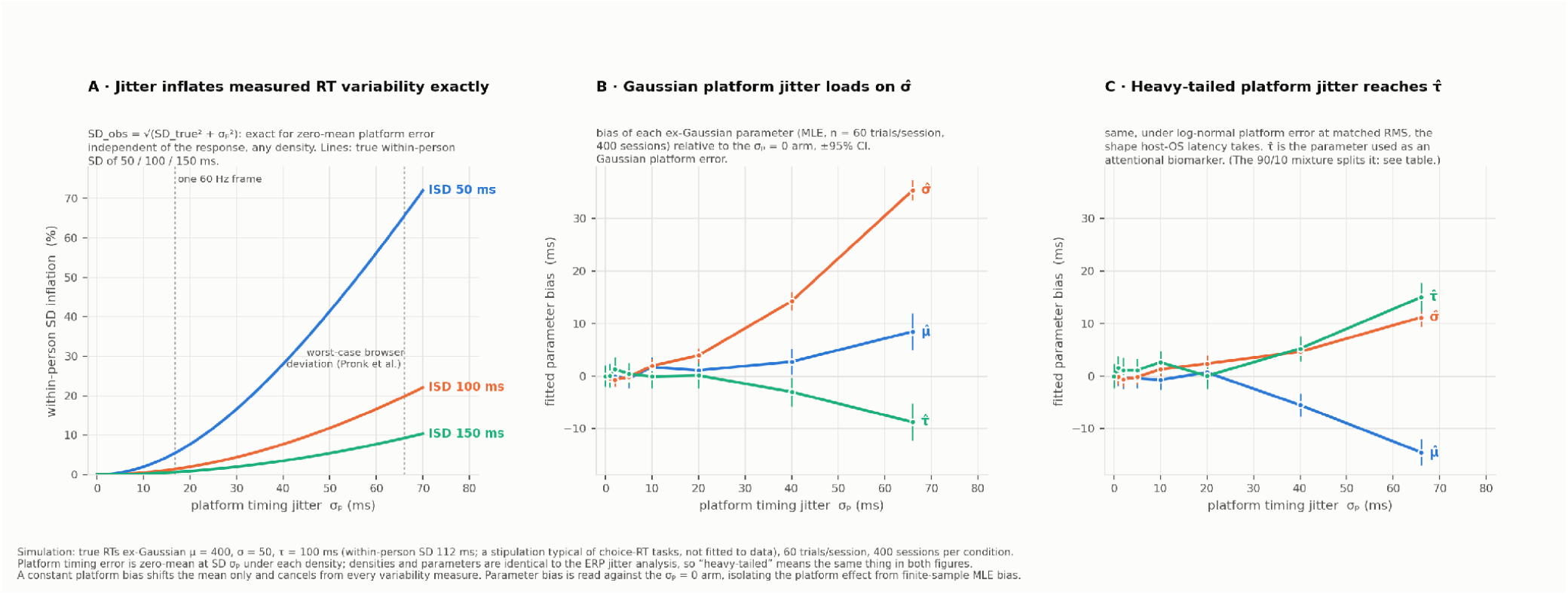
Platform timing variability propagates into within-person RT variability measures. (A)Zero-mean platform timing error of SD σ_p_, independent of the response, adds in variance. The observed SD is 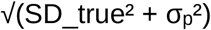, exactly and for any error density. Curves show the percent inflation of the measured within-person SD for true values of 50, 100, and 150 ms. Dashed references mark one 60 Hz display frame (16.7 ms) and the worst-case browser deviation of about 66 ms reported by Pronk et al. (2020). (B, C) Where the error lands in ex-Gaussian analyses. True RTs were simulated as ex-Gaussian with μ = 400, σ = 50, and 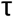= 100 ms. Each condition comprised 400 sessions of 60 trials. Platform error was added per trial and each session was refit by maximum likelihood. Bias is read against the σ_p_ = 0 arm of the same density, which isolates the platform effect from finite-sample fitting bias. Under Gaussian platform error (B), 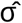 absorbs the error, gaining 35 ms at σ_p_ = 66 ms against a true σ of 50 ms, while 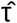 is pushed down by 9 ms. Under log-normal platform error at matched RMS (C), the error instead reaches 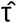, which gains 15 ms at σ_p_ = 66 ms. 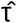 is the parameter used as an attentional biomarker. A 90/10 Gaussian-exponential mixture at the same RMS splits the error between 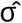 and 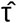 (supplementary table). All parameter biases at σ_p_ ≤ 5 ms are statistically indistinguishable from zero. The error densities and the true RT distribution are stipulations. The heavy-tailed shapes are not measurements of any specific platform.

LSL provides inter-stream clock synchronization, offset estimation, and cross-host alignment through a protocol documented and validated independently of any client application (Kothe et al., 2025). MERIT’s responsibility is the fidelity of the event timestamps it supplies to LSL. Four quantities define that fidelity. They are the offset, variability, device dependence, and within-session drift of one interval. That interval runs from a physical event on the participant device (photons appearing on the screen, a touch registering) to the LSL-clock timestamp MERIT assigns to it. The single-device precision reported above characterizes the variability of this interval but not its absolute offset. MERIT’s marker-to-photon and touch-to-marker offsets therefore remain to be characterized across representative devices and recording sessions. Until that work is complete, MERIT’s LSL support should be understood as an implemented integration pathway rather than a fully validated end-to-end timing solution. Presentation Mobile, the nearest native runtime in Table 1, supplies the benchmark. Woods et al. (2017) measured its event markers through LSL against an external timer and reported device-specific offsets of 1.3 to 23 ms with sub-millisecond variability. Per-device correction was required before performance could be compared across devices. Those measurements predate current devices, and the same characterization of MERIT on current-generation hardware is the next benchmarking step.

On the interval-precision dimension measured, MERIT’s Unity-based runtime performs substantially closer to dedicated laboratory hardware than to browser-based alternatives, with the residual variance attributable primarily to display hardware characteristics rather than to the platform itself. The four task families and the platform’s deployment infrastructure together support a range of research applications. We describe representative deployments that illustrate the platform’s use rather than provide exhaustive validation, which is the subject of separate reports.

### The cpCST Continuous Control Paradigm

cpCST illustrates the class of paradigm that the platform was designed to make broadly accessible. Prior continuous control paradigms have required laboratory-grade input hardware and dedicated software, limiting their use to a small number of laboratories with the technical capacity to support them. cpCST in MERIT runs on consumer devices, calibrates motor stability within approximately two minutes, and produces measures that have been validated against flanker performance, psychometric IQ, and VO2max (MacKay-Brandt et al., 2025). The paradigm is currently being deployed in studies of cognitive aging, Long COVID, and cardiovascular fitness as a modifiable risk factor for cognitive decline. Its delivery on participants’ personal devices through Curious-mediated EMA schedules enables longitudinal sampling at densities that laboratory administration could not support.

### Reaction-Time Paradigms in Laboratory and Remote Settings

Classical reaction-time paradigms (Stroop, flanker, n-back, go/no-go) have been built in MERIT and benchmarked against their established effects. The flanker congruency effect, for example, replicates with equivalent magnitude when the same task is administered through MERIT on a controlled laboratory desktop and on participants’ personal smartphones, consistent with the timing benchmarks reported above. The capacity to administer the same task instrument through the same engine across laboratory and remote settings supports a class of study designs (within-participant lab-to-field validation, large-scale remote replication of laboratory effects) that have previously required either accepting timing variability or running separate instruments.

Mean reaction-time effect sizes are the most familiar dependent measures in this family, but they are not the most demanding test of platform timing. Within-person reaction-time variability, intra-individual coefficient of variation, and the parameters of ex-Gaussian or other RT distribution fits are increasingly used as digital biomarkers of attentional regulation and early cognitive change, and these measures are highly sensitive to platform-induced jitter. A platform whose own timing variability is comparable to or larger than the within-person variability it is attempting to measure will systematically attenuate or distort these signals.

Variability-based measures make the timing requirement quantitative, because the arithmetic is exact. Zero-mean platform timing error of SD σ_p_ adds to the measured within-person variance, inflating the observed SD by the factor 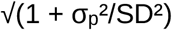 (Figure 4A). At millisecond-level jitter the contribution is negligible by any standard. Two milliseconds of jitter inflates a 100 ms within-person SD by 0.02%. At browser-class error the distortion is structural. The 66 ms worst-case deviation reported for smartphone browsers inflates a 50 ms within-person SD by two thirds. Distributional analyses fare worse, and in a way a single RMS figure cannot predict. In simulated ex-Gaussian fits (Figures 4B and 4C), Gaussian-shaped platform error loads on 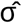 and pushes 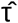 down. Log-normal error at the same RMS, the shape operating-system latency typically takes, instead raises 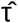 by 15 ms at the browser worst case. The tail shape of a platform’s timing error is a property of the device and operating system rather than of the participant. That shape determines whether the attentional biomarker is inflated or suppressed. Unlike sampling noise, the distortion is systematic. It does not average out over repeated sessions, and it shifts whenever a participant’s device or operating system changes mid-study. Those are precisely the conditions of longitudinal and burst-sampling deployment. The same arithmetic sets the stakes for individual-differences research. Mean reaction time is largely protected, because the variance of per-trial platform error is divided by the trial count when a session mean is formed. Variability-based measures have no such protection. When σ_p_ differs across participants’ devices, the platform adds between-person variance that has nothing to do with the participants, and that variance masquerades as an individual difference. A platform intended for variability-based digital biomarkers must therefore hold its timing variability an order of magnitude below the within-person SD under study. It must also report its measured timing distribution, including the tail, rather than a single summary statistic. MERIT’s frame-level timing logs exist to make that report possible per deployment. The benchmarks in the Performance Benchmarking section locate the platform’s σ_p_ on the axes of Figure 4.

### Digital Neuropsychological Drawing Tasks

The drawing family enables digital administration of paradigms with decades of normative data while extending those paradigms with kinematic measures. Clock drawing administered through MERIT, for example, preserves the traditional scoring criteria that clinicians rely on while also capturing per-stroke kinematic features of the kind prior work has used to help separate cognitive impairment from motor slowing; whether MERIT-captured kinematics support the same distinctions is an empirical question for dedicated validation studies. Similar extensions apply to the Rey-Osterrieth, digit-symbol substitution, and spiral drawing paradigms. These deployments are particularly suited to populations (older adults, clinical samples) for whom paper-and-pencil paradigms remain the familiar mode of administration and for whom abrupt transition to novel digital paradigms would compromise data comparability with existing literatures.

### Longitudinal and Burst-Sampling Designs

The integration of MERIT tasks with the Curious scheduling infrastructure supports EMA and burst-sampling designs in which participants complete brief task batteries repeatedly over weeks or months. In a recent deployment (The NKI Rockland Sample II: An Open Resource of Multimodal Brain, Physiology & Behavior Data from a Community Lifespan Sample, R01MH124045) within the NKI-Rockland Sample (Nooner et al., 2012), participants completed short cognitive batteries on their personal devices on a defined schedule, with the resulting data integrated with the cohort’s existing neuroimaging, physiological, and behavioral records.

The capacity to administer the same paradigm densely over time, on the same engine that delivered the in-laboratory baseline, supports inference about within-person cognitive dynamics that single-occasion or sparse repeated-measures designs cannot recover.

## Discussion

MERIT addresses a methodological gap that has constrained cognitive research as it has moved into real-world settings. Precise instruments have stayed in the laboratory, scalable instruments have run in the browser, and neither has carried the same task from synchronized laboratory recording to participants’ own devices. MERIT combines a no-code visual builder with a Unity-based runtime engine embedded natively within mobile host applications. Investigators without programming expertise can therefore deliver tasks with measured millisecond interval precision on the devices their participants carry. The same tasks stream to LSL in the laboratory and run on EMA schedules in the field.

Several implications follow from this design. The platform makes it feasible to administer the same task instrument in both laboratory and field settings, supporting a class of study designs (e.g., direct laboratory-to-field comparison, large-scale remote replication, longitudinal designs anchored by in-laboratory baselines) that the precision-accessibility trade-off has historically blocked. Investigators no longer need to choose between a controlled instrument that does not scale and a scalable instrument whose timing properties preclude rigorous inference.

More specifically, the Unity platform’s native LSL integration allows in-laboratory administration of MERIT tasks to be synchronized with EEG, fNIRS, eye-tracking, ECG, electrodermal activity, and other instrumented data streams. This integration supports an ecosystem of more than 150 supported device classes (Kothe et al., 2025), allowing characterizations of the brain and body state correlates of task performance that the same task in the field cannot directly measure. Field deployments of that same task, delivered through the same engine to participant devices, can then carry the interpretive weight of the laboratory characterizations: a reaction-time slowing observed in burst-sampling data can be interpreted against the cortical, autonomic, or attentional patterns established in the laboratory using the identical instrument. Laboratory recordings can thereby serve as the mechanistic grounding for ecological data, and ecological data extend the reach of laboratory findings into the contexts where cognition is actually deployed. The result is a single instrument that bridges what have historically been parallel and largely disconnected research literatures.

The platform also lowers the barrier to methodological innovation. Researchers with novel paradigm ideas can prototype, iterate, and deploy in a timeframe of days rather than months, and without recruiting a programmer or accepting the constraints of fixed task batteries. The four currently supported task families cover most existing paradigms, but the platform’s architecture is intended to grow with the field. Continuous control paradigms in particular, exemplified by cpCST, illustrate a class of measurement that the platform makes broadly accessible for the first time.

### Gamification and Engagement

Engagement is an underappreciated determinant of data quality in remote and longitudinal cognitive research. Participant compliance with repeated assessment schedules declines over weeks of EMA delivery, and within a session, sustained engagement directly shapes the validity of the cognitive measures themselves. Because MERIT is built on a high-performance interactive runtime, the platform supports a range of gamification affordances that have historically been impractical for cognitive paradigms: narrative framing, contingent visual and auditory feedback, scoring and progression structures, adaptive difficulty calibrated to individual performance, and aesthetic treatments that align tasks with the visual conventions participants encounter elsewhere on their devices. These affordances are applied through the builder rather than coded, and they layer onto the underlying cognitive paradigm without altering its measurement properties. The distinction is important: MERIT does not build games and does not transform cognitive paradigms into game-like structures whose construct validity has not been established. Instead, the platform makes gamification an instrument for engagement and compliance in the service of validated paradigms, with particular value for pediatric populations, repeated-measures designs, and remote deployments where participant attrition is a primary threat to inference.

### Open Science

MERIT is being developed within an open science framework. Task definitions are designed to be portable and shareable: investigators can publish their MERIT task specifications alongside their reports, allowing direct replication of the instrument rather than approximate reconstruction from method sections. This addresses a long-standing reproducibility concern in cognitive research, where two laboratories implementing “the same” paradigm from published descriptions can produce subtly different instruments. We are coordinating with the Open Measures Network Initiative (OMNI) on measurement harmonization standards and contributing the platform’s task taxonomy as a reference resource.

### Sustainability and the CMI Distribution Model

Long-term sustainability of open scientific infrastructure is a recognized challenge. Platforms that succeed in the research community frequently outlive the funding cycles that produced them, and the cost of supporting a production-grade platform across operating system updates, device populations, and evolving security requirements is non-trivial. MERIT is being distributed through the Child Mind Institute under a model analogous to Red Hat’s approach to open-source enterprise software: the platform itself is open and freely available for research use, while CMI provides supported deployment, hosting, and integration services to laboratories, consortia, and clinical operations that require service-level guarantees. This structure aligns the platform’s open availability with a sustainable funding base for ongoing engineering and support, and it positions the platform to remain available to the broader research community independent of any single grant cycle.

### Limitations, Ongoing and Future Development

Several limitations bear acknowledgment. Absolute timing on mobile devices, while substantially improved relative to browser-based alternatives, remains influenced by operating system constraints (background processes, battery optimization features, vendor-specific scheduling) that can introduce occasional frame-level deviations. We are profiling these effects across a broader device population and developing calibration routines that run briefly on each participant device before task execution, allowing investigators to characterize and account for device-specific timing properties. Display rise and fall times remain a hardware-level consideration that no software platform can eliminate; we report measured values for representative devices so that investigators can make informed decisions for paradigms sensitive to onset definition.

Second, integration with the Curious deployment infrastructure means that adopting MERIT involves adopting the Curious participant-facing application as the delivery channel. We are evaluating additional deployment paths, including standalone-application export and integration with other participant-facing platforms, to support investigators whose existing infrastructure assumes a different delivery model.

Third, the platform’s current task families, while broad, are not exhaustive. Virtual reality paradigms, paradigms requiring specialized input hardware, and paradigms that depend on continuous external data streams (beyond LSL-supported devices) are areas of ongoing development, as is the dynamic adaptive testing described under Task Families.

## Conclusion

MERIT unifies millisecond-precise mobile cognitive assessment, laboratory neurophysiological integration, and longitudinal remote measurement within a single no-code ecosystem. Investigators without programming expertise can design and deploy cognitive tasks with measured millisecond interval precision on consumer devices, in both controlled laboratory and naturalistic field settings. Because the same instrument can run in the laboratory alongside synchronized EEG, fNIRS, and physiological recordings and at home on participants’ own devices, behavioral signatures observed during ecological and burst-sampling deployments can be grounded in the brain and body state correlates established in laboratory administrations of the identical task. The platform’s four current task families cover most contemporary cognitive assessment paradigms. Its Unity-based runtime and native mobile embedding deliver measured interval precision an order of magnitude below the most demanding requirement estimated here, and its native LSL integration provides the path to synchronized multimodal recording. Its open distribution through the Child Mind Institute provides a sustainable basis for long-term availability to the research community. The platform’s broader contribution is to relax a trade-off that has structurally limited cognitive research as it has attempted to engage with real-world environments, making it feasible to administer the same instrument, with the same precision and the same interpretive backing, across the full range of settings in which cognition occurs.

## Methods

### Timing requirements for ERP-synchronized paradigms

We estimated timing requirements from ERP CORE (Kappenman et al., 2021), an open compendium of ERP paradigms recorded from 40 adults aged 18 to 30. We analyzed the BIDS release of the raw continuous recordings. Data were acquired with a BioSemi ActiveTwo amplifier at 1024 Hz, with 30 scalp channels, 3 periocular channels, and no online filtering. Three components spanning the spectral range of common ERPs were analyzed: the N170 at PO8, the MMN at FCz, and the P3 at Pz. The ERP analysis is a secondary analysis of the publicly available, de-identified ERP CORE dataset, and required no additional ethical approval.

Periocular channels were combined into bipolar HEOG and VEOG derivations. Continuous data were DC-corrected and high-pass filtered at 0.1 Hz with a zero-phase Butterworth filter. Ocular artifacts were removed with extended-infomax ICA. Components matching the bipolar EOG derivations were excluded automatically, between one and three per recording. Referencing, epoching, baseline correction, artifact rejection, measurement electrodes, and measurement windows followed the published ERP CORE parameters (Kappenman et al., 2021, Tables 1 and 2). Event codes for the two visual components were corrected for the recording monitor’s display delay. The dataset applies this delay but does not distribute its value. We measured it from the data as 17.81 ms, jointly across the two visual components. Usable samples were 39 participants for the N170, 38 for the MMN, and 39 for the P3.

Before any jitter condition was run, the pipeline at zero jitter was required to reproduce the component amplitudes published for this dataset (Kappenman et al., 2021, Table 3). No jittered result was read before this gate passed.

Latency jitter was then injected per trial as an exact fractional-sample shift, implemented as a phase ramp in the Fourier domain with 250 ms of zero padding. The jitter SD σ took values of 0, 1, 2, 5, 10, and 20 ms. Three error densities were used: a Gaussian, a log-normal with shape parameter 0.75, and a mixture of a 90% Gaussian core with a 10% exponential tail. The mixture’s tail scale was set so the tail carries half the total variance. Each density was affinely mapped to zero mean and SD σ. Any departure from the Gaussian prediction is therefore attributable to tail shape alone. Both heavy-tailed densities are stipulations rather than measurements of any specific platform. Ten independent jitter realizations were drawn per participant per condition. An independent prediction of each degraded average was computed by multiplying the spectrum of the clean average by the characteristic function of the jitter density. This prediction uses no information from the jittered trials.

The amplitude estimand was the mean amplitude of the condition difference wave in the published window at the published electrode. The latency estimand was the peak latency within the published search window. For each measure we set a materiality threshold of half its between-participant SD. The tolerable-jitter ceiling σ* is the smallest σ at which the worse of the two biases crosses its threshold. Because the full bias curves are reported (Figure 2C), the ceiling under any alternative criterion can be recovered from those curves. Jitter was modeled as independent across trials. Temporally correlated timing error and discrete frame-drop distributions, which deployed platforms can produce, are not covered by this model.

All analysis parameters were fixed and committed to version control before any EEG sample was read. Amendments were committed before the runs they affected. Analyses used Python 3.12 with MNE 1.12.1, MNE-BIDS 0.15.0, NumPy 2.2.6, and SciPy 1.18.0. The full pipeline is provided in the supplement (see Data and code availability).

### Simulation of platform jitter and RT variability measures

No human data were used in this analysis. True reaction times were drawn from an ex-Gaussian distribution with μ = 400 ms, σ = 50 ms, and τ = 100 ms. These values are typical of choice-RT tasks and are stipulations rather than fits to data. The implied within-person SD is 112 ms. Each simulated session comprised 60 trials, and each condition comprised 400 sessions.

Platform timing error was added to each trial as an independent zero-mean draw with SD σ□, for σ□ of 0, 1, 2, 5, 10, 20, 40, and 66 ms. The three error densities and their parameters were identical to those of the ERP analysis, so the two figures share one definition of heavy-tailed error. A constant platform bias shifts the mean only and cancels from every variability measure, so it was omitted. The model also assumes platform error is independent of the true response. Input-scanning latencies that correlate with response phase would add structure this simulation does not capture.

For each session we computed the sample SD and fit an ex-Gaussian by maximum likelihood (SciPy exponnorm) with moment-based starting values. Parameter bias was read against the σ_p_ = 0 arm of the same density. This isolates the platform effect from finite-sample fitting bias, which affects both arms equally. The analytic prediction for SD inflation, observed 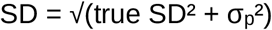, is exact and was plotted directly. The simulation used a fixed seed and is fully reproducible from the included script. It ran on Python 3.10 with NumPy 1.26.4 and SciPy 1.12.0.

## Data and code availability

ERP CORE is openly available (doi 10.18115/D5JW4R); we redistribute none of its raw data. The code and group-level results behind the timing-requirements analysis and the RT-variability simulation are available at https://github.com/SympatiCog/merit-timing-supplement(commit 79f522c).

## Acknowledgments

MERIT and Curious part of a broader portfolio of scalable digital mental health tools developed by the Child Mind Institute. We thank Metalab and ScienceSoft for their contributions to the development of the Curious and Mirror platforms. The development and dissemination of MERIT and the broader Curious platform have been supported in part through the Child Mind Institute’s partnership with the California Department of Health Care Services under the Children and Youth Behavioral Health Initiative, as well as by the Stavros Niarchos Foundation. Additional support provided by R01MH124045 and R01MH130901. MPM is the Phyllis Green and Randolph Cowen Scholar at the Child Mind Institute.

